# Comparative analysis of serological assays and sero-surveillance for SARS-CoV-2 exposure in US cattle

**DOI:** 10.1101/2024.04.03.587933

**Authors:** Santhamani Ramasamy, Meysoon Qureshi, Swastidipa Mukherjee, Sonalika Mahajan, Lindsey Cecelia LaBella, Shubhada Chothe, Padmaja Jakka, Abhinay Gontu, Sougat Misra, Meera Surendran-Nair, Ruth H. Nissly, Suresh V. Kuchipudi

## Abstract

Coronavirus disease-2019 (COVID-19) caused by severe acute respiratory syndrome coronavirus-2 (SARS-CoV-2) continues to pose a significant threat to public health globally. Notably, SARS-CoV-2 demonstrates a unique capacity to infect various non-human animal species, documented in captive and free-living animals. However, experimental studies revealed low susceptibility of domestic cattle (*Bos taurus*) to ancestral B.1 lineage SARS-CoV-2 infection, with limited viral replication and seroconversion. Despite the emergence of viral variants with potentially altered host tropism, recent experimental findings indicate greater permissiveness of cattle to SARS-CoV-2 Delta variant infection compared to other variants, though with limited seroconversion and no clear evidence of transmission. While some studies detected SARS-CoV-2 antibodies in cattle in Italy and Germany, there is no evidence of natural SARS-CoV-2 infection in cattle from the United States or elsewhere. Since serological tests have inherent problems of false positives and negatives, we conducted a comprehensive assessment of multiple serological assays on over 600 cattle serum samples, including pre-pandemic and pandemic cattle sera. We found that SARS-CoV-2 pseudovirus neutralization assays with a luciferase reporter system can produce false positive results, and care must be taken to interpret serological diagnosis using these assays. We found no serological evidence of natural SARS-CoV-2 infection or transmission among cattle in the USA. Hence, it is critical to develop more reliable serological assays tailored to accurately detect SARS-CoV-2 antibodies in cattle populations and rigorously evaluate diagnostic tools. This study underscores the importance of robust evaluation when employing serological assays for SARS-CoV-2 detection in cattle populations.

## Introduction

Coronavirus disease-2019 (COVID-19), caused by the severe acute respiratory syndrome coronavirus-2 (SARS-CoV-2), will remain a threat to public health for the foreseeable future. A remarkable feature of SARS-CoV-2 is the ability to infect many non-human animal species, and natural SARS-CoV-2 infection of multiple captive and free-living animals has been documented[1-6]. Receptor binding and membrane fusion are critical steps for coronaviruses to cross the species barrier and establish efficient transmission pathways in new host species. SARS-CoV-2 spike (S) protein mediates virus entry and cell fusion through its direct interaction(s) with the cellular angiotensin-converting enzyme-2 (ACE2) receptor[7-9]. The ability of the S protein to bind to ACE-2 receptors is a crucial determinant of host susceptibility to SARS-CoV-2 infection.

Comparative and structural analysis of ACE2 receptors in vertebrates predicted that several mammals could be at high risk for SARS-CoV-2 infection[8-10]. Based on ACE2 binding to the receptor binding domain (RBD) of the S protein of wildtype B.1 lineage, domestic cattle (*Bos taurus*) have been predicted to be susceptible to SARS-CoV-2[2]. Subsequently, experimental studies showed low susceptibility of cattle to experimental ancestral B.1 lineage SARS-CoV-2 infection with low levels of viral replication and limited seroconversion[11,12]. SARS-CoV-2 continues to evolve, resulting in the emergence of mutational variants named after the Greek letters Alpha, Beta, Gamma, Delta and Omicron. The emergence of variants might result in altered host tropism. For example, laboratory mice that were resistant to wildtype SARS-CoV-2 infection were found to be susceptible to alpha and other variants[13-17]. More recently, experimental co-infection of calves found that cattle are more permissive to infection with SARS-CoV-2 Delta than Omicron BA.2 and Wuhan-like isolates[18] . Further, the study also found limited seroconversion and no clear evidence of transmission to sentinel calves[18]. A study in 2022 reported detection of SARS-CoV-2 antibodies in lactating cows in Italy[19]. Subsequently, a serological survey in Germany found antibody evidence of natural SARS-CoV-2 exposure of cattle[20]. While these studies raise concerns about the potential spillover of recent SARS-CoV-2 variants into cattle, there is currently no evidence of natural SARS-CoV-2 infection in cattle from the United States or elsewhere in the world.

Given the experimental evidence indicating low susceptibility, limited seroconversion, and a lack of horizontal transmission of SARS-CoV-2 among cattle, coupled with concerns about the specificity of serological assays, it becomes imperative to thoroughly evaluate various serological methods for detecting SARS-CoV-2-specific antibodies in cattle. Consequently, we conducted a comprehensive assessment using multiple serological assays on over 600 cattle serum samples, including pre-pandemic and pandemic sera. We found no serological evidence of natural SARS-CoV-2 infection and transmission of SARS-CoV-2 in cattle in the USA. This study emphasizes the importance of rigorous evaluation when employing serological assays for SARS-CoV-2 detection in cattle populations.

## Materials and Methods

The following materials were obtained through BEI Resources, NIAID, NIH: human embryonic kidney cell line expressing human angiotensin-converting enzyme 2 (HEK-293T-hACE2) (NR-52511); SARS-Related Coronavirus 2 Wuhan-Hu-1 Spike-Pseudotyped Lentiviral Kit V2, (NR-53816). Plasmids encoding spikes of SARS-CoV-2 variants Delta (Cat. No. 172320) and Omicron (Cat. No. 179907) were procured from Addgene, USA.

### Serum samples

Serum collected from cattle (n=549) from early 2022 to 2023 for the screening of bovine viral diseases at Animal Diagnostic Laboratory at Pennsylvania State University were analyzed in this study for the presence of SARS-CoV-2 antibodies. The age of cattle tested ranged from 2 months and older. Cattle sera (n=49) collected before 2020 were used as pre-pandemic negative controls. Additionally, hyperimmune sera (n=3) from cattle immunized with B.1 lineage RBD protein described in our earlier publication were included as positive controls in some assays. All animal care and sample collections were approved and performed in accordance with the guidelines of the Institutional Animal Care and Use Committee at Pennsylvania State University. The Pennsylvania State University Institutional Animal Care and Use Committee (IACUC protocol # PROTO202001506).

### Production of SARS-CoV-2 pseudoviruses

SARS-CoV-2 spike pseudoviruses were produced using the third-generation lentiviral plasmids as described elsewhere [21]. Lentiviral helper plasmid encoding Gag/pol, transfer plasmid encoding luciferase and ZsGreen, Tat and Rev and plasmid encoding spike of SARS-CoV-2 variants Delta or Omicron were transfected in HEK 293T cells using Fugene6 reagent (Cat. No. E2691, Promega) following manufacturer’s guidelines. The pseudovirus containing cell culture supernatants were collected after 48 hours of transfection, and filtered aliquots were stored at - 80°C until use. The infectivity of SARS-CoV-2 pseudoviruses were determined using HEK-293T-hACE2 cells. Briefly, the HEK-293T-hACE2 cells were infected with 10-fold serial dilutions of pseudoviruses in 96 well clear bottom plate (Cat. No. 165306, ThermoScientific, USA). At 72 hours post infection, RLUs were measured (BioTek Synergy HTX Multi-Mode Microplate Reader, Agilent) following the addition of BrightGlo luciferase reagent (Cat. No. E2620, Promega). The dilution of the virus that showed ∼10^4^ RLUs was used in the pseudovirus neutralization assay.

### Detection of SARS-CoV-2 antibodies using pseudovirus neutralization assay (pVNT)

We employed pVNT to test the presence of SARS-CoV-2 neutralizing antibodies in cattle sera using pseudoviruses containing spike proteins from Delta and Omicron SARS-CoV-2 variants of concern (VoCs). Briefly, the pseudoviruses equivalent of 10^4^ RLUs were incubated with 1:30 dilutions of heat inactivated sera for an hour at 37 °C. The pseudovirus/sera mixtures were inoculated into 96-well plates containing 1.3 × 10^4^ HEK-293T-hACE2cells. The pseudovirus infectivity was determined at 72 hours post infection by quantifying the luciferase activity. The percentage neutralization of pseudoviruses was calculated by normalization to a virus-only control. Each serum was tested in a single well initially and the samples with percent neutralization of ≥60% were further tested in duplicates at three dilutions (1:30, 1:60, 1:120 and 1:240). A percent neutralization of 60% was further tested in other assays. The results were analyzed using GraphPad Prism Software version 9 (San Diego, CA, USA).

### Live Virus neutralization (VN) assay for SARS-CoV-2 antibodies

VN assays to determine SARS-CoV-2 neutralizing antibody titers were performed as described earlier[22]. Briefly, Vero E6 cells were seeded onto 96-well plates and cultured for 18-24 hours at 37°C with 5% CO_2_. Serum samples were heat inactivated, diluted 2-fold in triplicates using DMEM and mixed with 100TCID_50_ of SARS-CoV-2 [hCoV-19/USA/PHC658/2021 (lineage B.1.617.2; Delta), and hCoV-19/USA/MD-HP20874/2021 (lineage B.1.1.529; Omicron)] and incubated at 37°C for one hour. The serum-virus mixtures were added to Vero E6 culture and incubated for 72 hours. Cells were observed for cytopathic effects under an inverted light microscope. The reciprocal of the highest dilution of serum showing no cytopathic effects in at least two out of three wells is considered as the neutralization titer of the serum.

### Live virus neutralization assay for Bovine coronavirus antibodies

We performed a virus neutralization assay to detect bovine coronavirus specific antibodies following a previously reported procedure[23]. The two-fold serial dilutions of heat inactivated serum were mixed with 100 TCID_50_ of bovine coronavirus strain Mebus and incubated for one hour at 37 °C with 5% CO_2_. The virus and serum mixture were added to MDBK cells, grown in a 96-well microtiter plate, and incubated for 4 to 5 days at 37 °C with 5% CO_2_. The assay was performed in quadruplicate and endpoint neutralization titer was designated as the reciprocal of highest serum dilution, at which the virus infection is inhibited in all 3, or 2 of 3 wells as assessed by visual examination.

### SARS-CoV-2 Surrogate virus neutralization assay (sVNT)

We used the widely accepted species-agnostic SARS-CoV-2 antibody detection kit, GenScript cPass™ technology-based neutralization assay[24] for testing the cattle serum samples. The sVNT is useful for the detection of SARS-CoV-2 specific antibodies in human and animal species. We tested pandemic and prepandemic serum samples in SARS-CoV-2 Delta and Omicron based sVNTs using manufacturer’s instructions. Briefly, the serum samples were incubated with horse radish peroxidase (HRPO)-conjugated RBD (Delta or Omicron) (Cat. No. Z03614-20 and Cat. No. Z03730-20) and added to the wells coated with human ACE2 protein. Each serum was tested in single wells. The interaction of HRPO-conjugated RBD and ACE2 were determined by measuring the absorbance values after adding the developing solution. The wells showing >30% of inhibition was determined as positive for the antibodies.

### Indirect ELISA assay for SARS-CoV-2 antibody detection

We employed in-house developed indirect ELISA for the detection of antibodies in cattle serum samples [25]. Briefly, SARS-CoV-2 RBD antigens expressed in 293T cells were used to coat the 96-well ELISA plates (Cat No. 44240421, Thermofisher, USA). 50µL (2µg/mL) of antigens were added on the wells and incubated at 4°C overnight. Plates were washed thrice with PBS containing 0.05% Tween20 and blocked using 200 µL/well of Stabilguard immunoassay buffer (SG01-1000, Surmodics, MN, USA). After washing, the plates were incubated with the serum samples diluted in Stabilguard buffer (1:50) for one hour at 37°C. Then 100µL of anti-bovine IgG peroxidase (Cat # A5295, Sigma-Aldrich, MO, USA) was added to the wells at 1:10,000 dilutions. Plates were washed and incubated with 100µL per well of substrate containing 3,3′,5,5′-Tetramethylbenzidine dihydrochloride (Cat # T3405, Sigma-Aldrich, MO, USA) and hydrogen peroxide for 10 minutes. The reactions were terminated using 3N HCl and OD values were measured at 450nm using Cytation5 multi-mode reader. The samples showing OD values higher than the cut-off values were determined as positive for SARS-CoV-2 antibodies.

## Results

### Pseudovirus neutralization assay suggests SARS-CoV-2-specific antibodies in cattle serum of varying quality

In total 549 pandemic serum samples and 49 pre-pandemic serum samples were tested in SARS-CoV-2 pseudovirus neutralization assays (pVNT). We have previously demonstrated high cross-reactivity of ancestral B.1 RBD-specific hyperimmune serum against pseudovirus expressing pre-Omicron variant spike protein but low cross-reactivity against Omicron pseudovirus [26]. Therefore, pVNT using Delta (pre-Omicron) and Omicron pseudoviruses were performed. Out of 549 pandemic samples, 56 serum samples showed >60% inhibition in pVNT using Delta, and 44 serum samples had >60% inhibition in pVNT using Omicron pseudoviruses. The sixty percent inhibition indicates that the percent inhibition at serum dilution 1:30. However, none of the samples showed >90% inhibition at the 1:30 dilution of serum. Therefore, the 50% neutralization titer lies around 30 which is a very low or inconclusive titer. Note that 60% inhibition in pVNT is not a positive-negative cut-off in pVNT **(Figure 1)**. Interestingly, two of the 49 pre-pandemic serum samples had >60% inhibition in pVNT using Delta spike. The quality of serum samples tested were variable, from pale and clear to red or dark brown with debris from blood. To rule out the effect of hemolysis on pVNT results, 33 pale and clear sera and 24 hemolyzed sera were randomly selected for the comparison of percent inhibition in pVNT. Three-fold serial dilutions (1:30 to 1:240) of the samples were tested in pVNT. In pVNT, 33% and 9% of pale/clear and 20% and 16% of hemolyzed samples showed >60% inhibition of RLUs at 1:30 dilution with Delta and Omicron spike pseudoviruses, respectively. Hemolysis and serum quality did not significantly impact whether specimens were above or below 60% inhibition, per two-sided Fisher’s exact test (delta *p*=0.56; omicron *p*=0.13).

**Figure 1:**
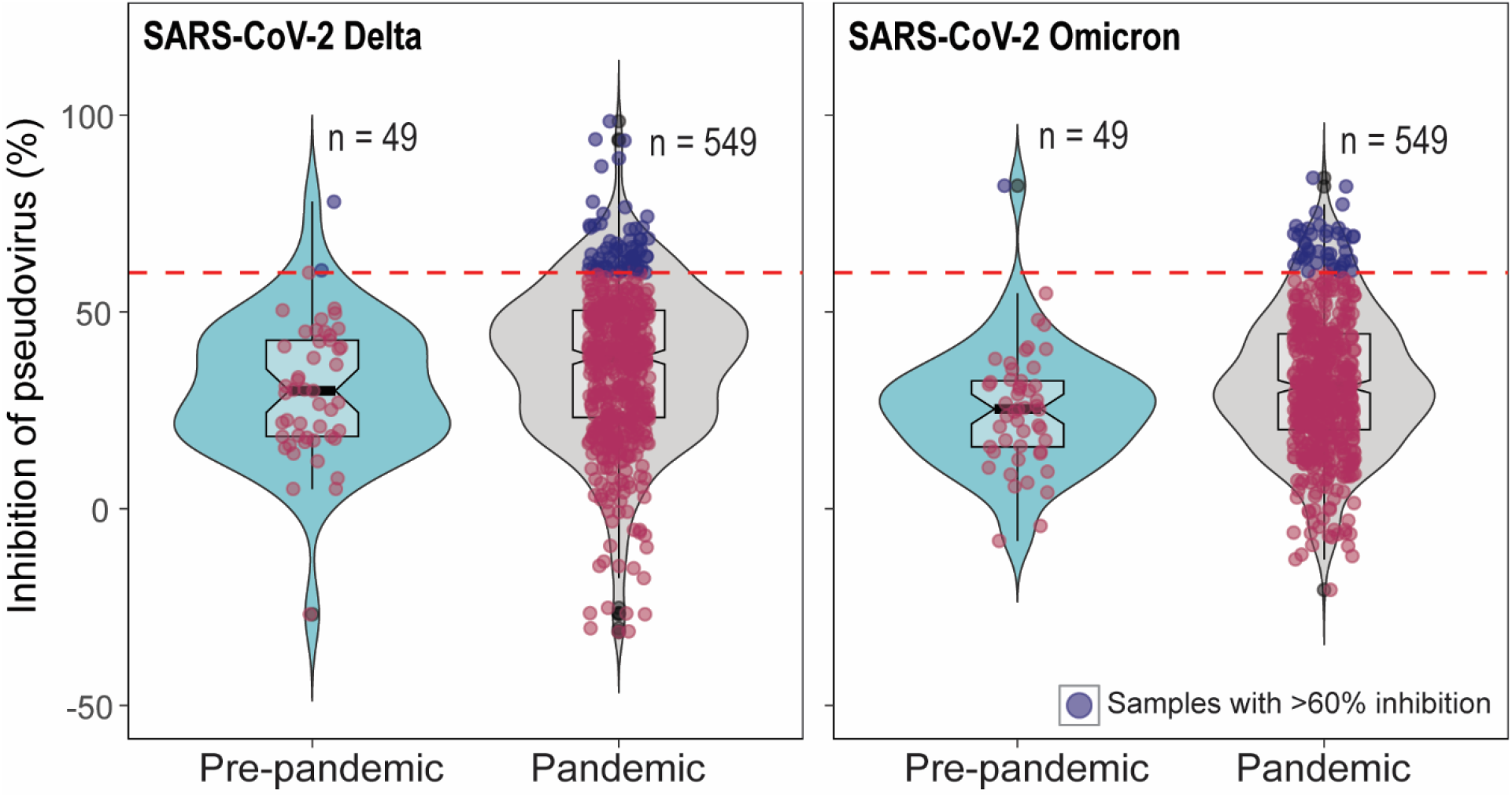
Distribution of percent inhibition of SARS-CoV-2 Delta and Omicron spike pseudoviruses by cattle serum samples in pVNT. In the pVNT, 549 pandemic and 49 pre-pandemic serum samples were tested. The dotted line indicates the 60% inhibition. 69 pandemic and two pre-pandemic serum showed >60% inhibition in SARS-CoV-2 Delta pVNT and 44 pandemic serum samples and one pre-pandemic serum showed >60% inhibition in Omicron pVNT.

### High percent inhibition in pVNT does not correspond to positivity in sVNT, indirect ELISA and VN

To confirm whether samples with pseudovirus inhibition indicated presence of SARS-CoV-2-specific antibody, we further tested the serum samples with >60% inhibition in pVNT using two additional assays measuring antibody binding to SARS-CoV-2 RBD. First, we tested sera in surrogate virus neutralization tests (sVNT) using RBD from Delta and Omicron [24,26,27]. Out of 90 samples (52 samples with >60% inhibition and 38 pre-pandemic samples), only two showed the percent inhibition above the cut-off in Delta sVNT. Of the 92 samples tested in Omicron sVNT, one sample showed the percent inhibition just above the cut-off **(Figure 2)**. The cattle that showed 55% Delta sVNT inhibition had 71.5% Delta pVNT inhibition; on the other hand, the serum with 33% Delta sVNT inhibition had 4% inhibition Delta in pVNT. The serum with 31% Omicron sVNT inhibition showed 59.5% inhibition in Omicron pVNT.

**Figure 2.**
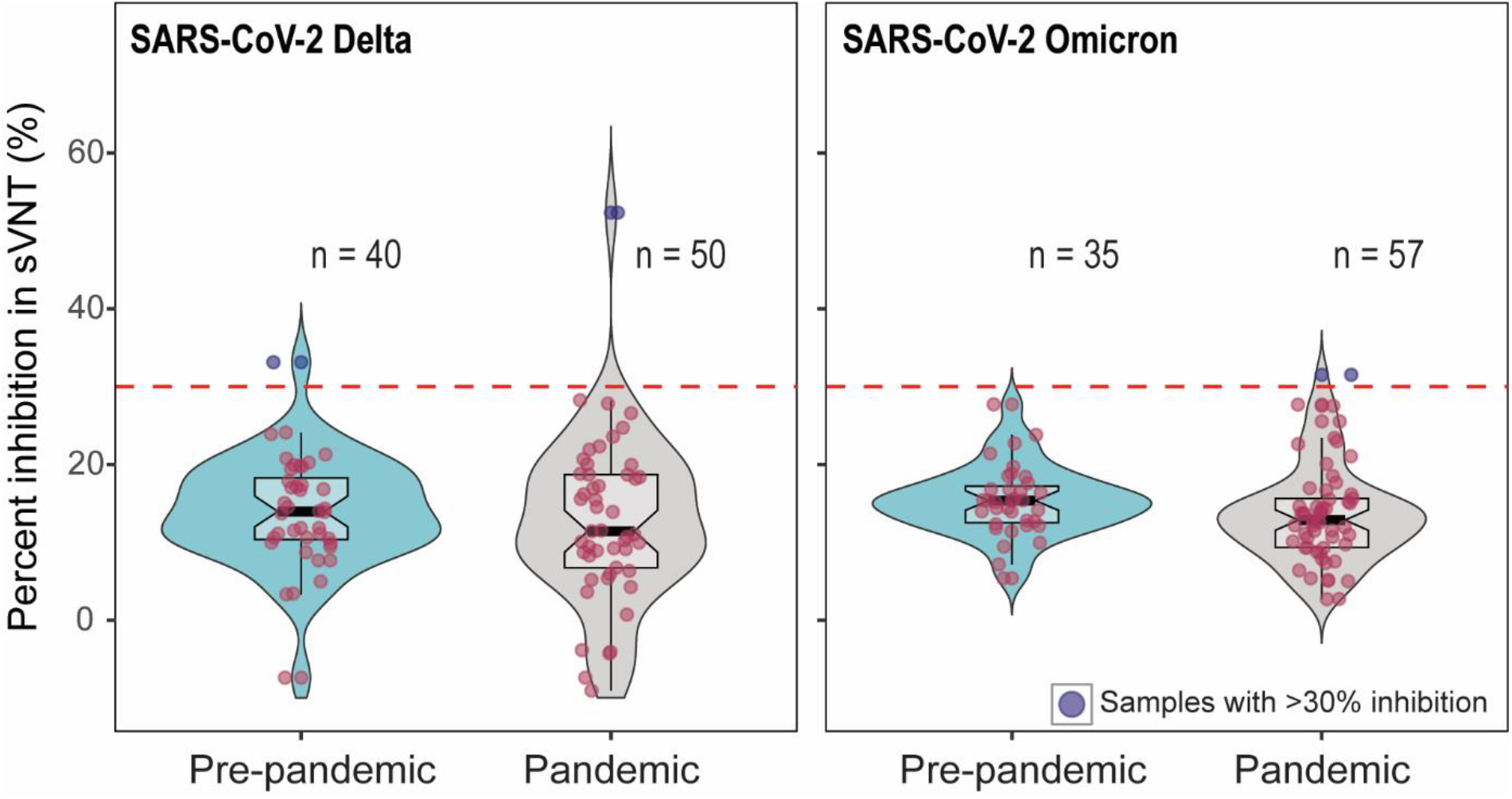
The percent inhibition of cattle sera in Delta (a) and Omicron (b) -RBD based sVNT. The positive-negative threshold stated by the manufacturer is 30%. Two out of 90 and one out of 92 serum samples showing >30% inhibition in Delta and Omicron sVNT, respectively.

We previously validated an ancestral B.1 lineage RBD indirect ELISA assay with 100% sensitivity and specificity compared to a live virus neutralization assay[25]. When serum samples (n=88) that showed >60% inhibition in pVNT were tested in this assay, one sample showed absorbance above the determined cut-off and 87 samples had absorbance below the cut-off **(Figure 3)**. Further, the samples that showed >30% inhibition in Delta (n=2) and Omicron (n=1) sVNT were negative in the indirect ELISA assay. The serum (n=1) that was positive in indirect ELISA had 45% inhibition in Delta pVNT. The serum samples with positivity in at least one of the serological assays are indicated in **Table 1**. The serum samples with >60% inhibition in pVNT and pre-pandemic samples were tested in live virus neutralization assays; none of the samples showed the neutralization at 1:20 dilution.

**Table 1.**
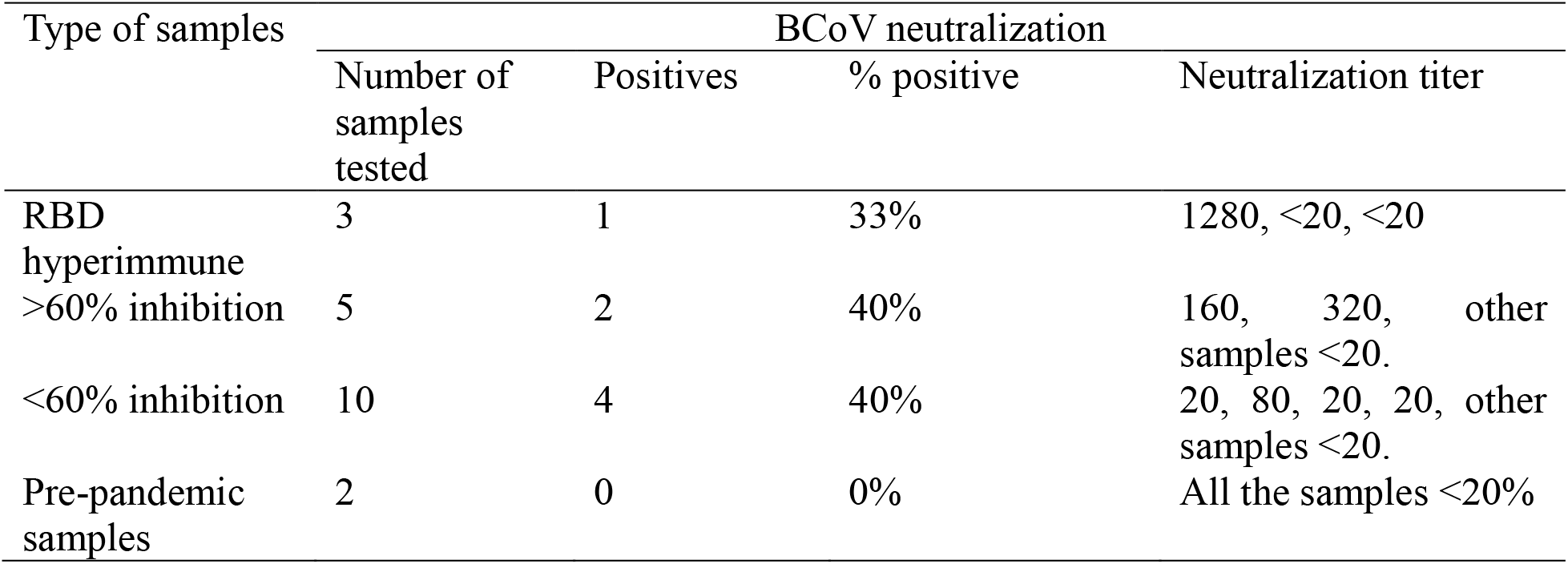
Determination of Bovine coronavirus neutralization titer of pre-pandemic (n=2), pandemic (n=15) and hyperimmune (n=3) serum samples.

**Figure 3.**
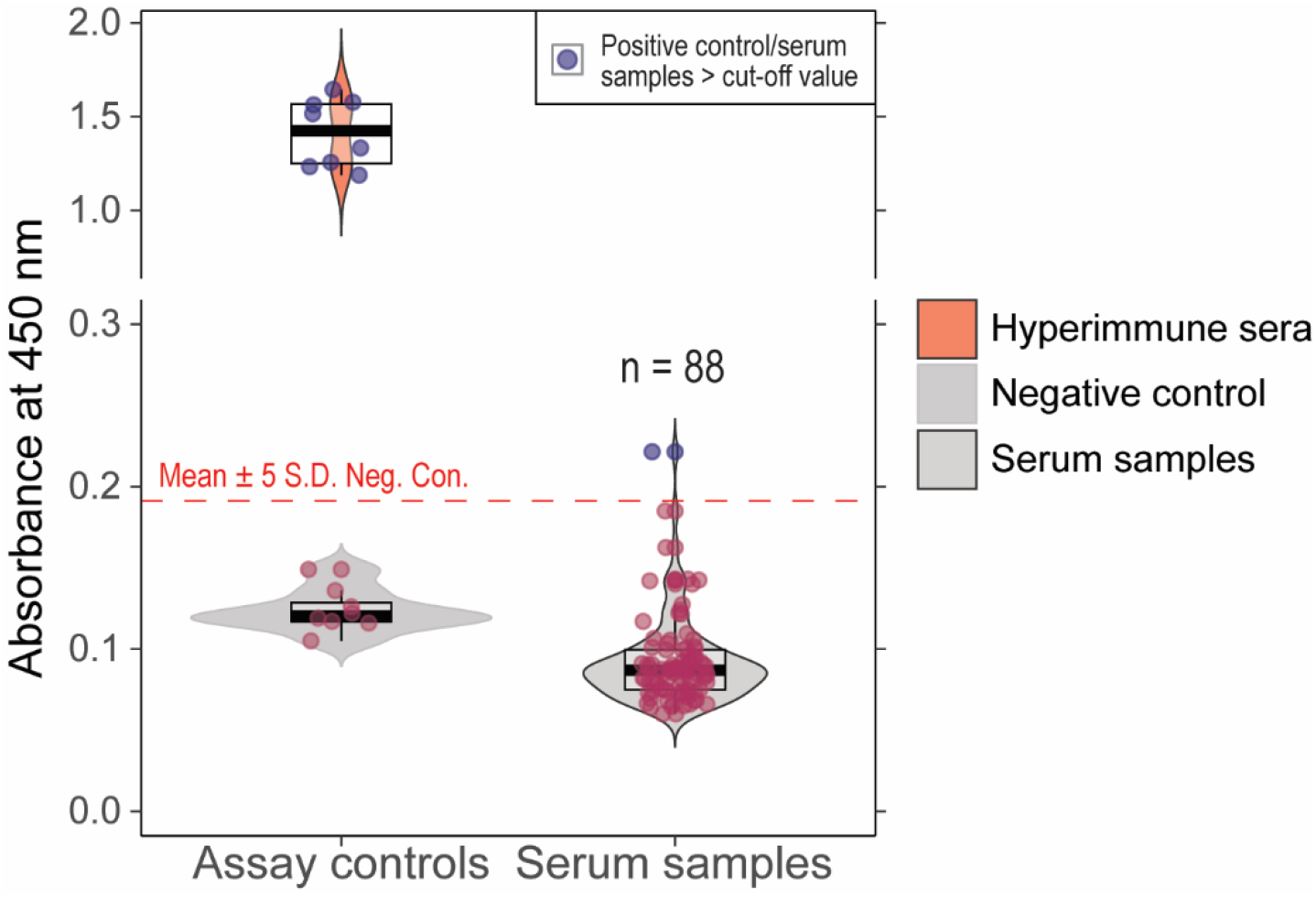
Absorbance values (A_450_ nm) for cattle serum samples tested in in-house developed indirect ELISA. The cut-off for the positive vs. negative samples is Mean+5S.D. One of the serum samples had absorbance values higher than the cut-off.

### SARS-CoV-2 specific cattle antibodies are not cross-reactive to Bovine coronavirus

Bovine coronavirus (BCoV), like SARS-CoV-2, is a member of the *Betacoronavirus* genus. BCoV is widespread in cattle populations, a host in which it can cause respiratory and enteric infections. Vaccination against BCoV is a common management strategy in the US. To understand if our observed SARS-CoV-2 pseudovirus inhibition could be due to cross-reactive bovine coronavirus antibodies, we tested a subset of cattle serum samples in BCoV live virus neutralization assays. We analyzed 3 hyperimmune sera raised in cattle against SARS-CoV-2 Wuhan RBD, 5 serum samples that showed >60% inhibition in pVNT, 10 samples that showed <60% inhibition in pVNT, and 3 prepandemic serum samples (a serum showed >60% inhibition in Omicron pVNT). One hyperimmune serum, two pandemic serum samples with >60% inhibition, and four pandemic serum samples with <60% inhibition in pVNT showed neutralization of BCoV (**Table 2**). The majority of samples (60%) demonstrated no neutralization of BCoV irrespective of pVNT status. These results indicate that the percent inhibition in SARS-CoV-2 pVNT is not in correlation with BCoV neutralization. Indeed, SARS-CoV-2 Wuhan RBD hyperimmune sera that had > 90% inhibition in pVNT demonstrated no cross-neutralization of BCoV (**Table 2**).

**Table 2:** Results demonstrating “positivity” in at least one SARS-CoV-2 serological assay.

| Serum id | Delta<br>pVNT | Omicron<br>pVNT | Delta<br>sVNT | Omicron<br>sVNT | Indirect<br>ELISA |
| --- | --- | --- | --- | --- | --- |
| P2231751-2 | 71% | 84% | 52% | Neg | Neg |
| P2002039-2 | 4% | 20% | 33% | Neg | Neg |
| 2C | 49% | 59.5% | Neg | 31% | Neg |
| P2214655-57 | 58.3% | 64% | Not done | Neg | Pos |

## Discussion

Cross-host transmission can occur with contact between viruses and potential new hosts [28]. Potential sources of exposure of cattle to SARS-CoV-2 include infected humans and animals. Abundant, sustained, and protracted human-to-human transmission of SARS-CoV-2 promotes the risk of spillover to susceptible animal species. Natural infection and circulation of SARS-CoV-2 has been well-established in white-tailed deer, the most abundant large mammal species in the US[29-33]. Shared home ranges enhance the potential for spillover of SARS-CoV-2 from white tailed-deer to cattle. It is well-established that bacterial and viral pathogens can be transmitted between deer and cattle due to the overlap of deer home ranges with cattle pastures. *Mycobacterium tuberculosis* and bovine viral diarrhea viruses are thought to persist through bidirectional transmission between cattle and deer[34,35], and transmission between species has been documented for other non-vector-borne pathogens including hepatitis E virus[36] and bovine coronavirus[37]. Transmission of SARS-CoV-2 within the human population occurs through multiple methods, including aerosols, droplets, and fomites, with spreading possibly through either direct or indirect contacts[38,39]. The potential for transmission of SARS-CoV-2 to livestock from wildlife through similar routes is high.

Earlier studies suggested cattle are poorly permissive to infection with SARS-CoV-2[11,12]. Further, a recent study suggested that cattle are more permissive to infection with SARS-CoV-2 Delta than Omicron BA.2[18]. Further, the study also found limited seroconversion and no clear evidence of transmission to sentinel calves[18]. Serological studies from Italy[19] and Germany[20] reported antibody evidence of natural SARS-CoV-2 exposure of cattle. Serological assays show various sensitivity ranges[40,41], and false-positive serology test results have been reported in COVID-19[42,43]; therefore, it is crucial to compare various serological testing methods in a given host species for serological determination of SARS-CoV-2 infection.

We investigated antibody presence in cattle using an easily adaptable pseudovirus neutralization assay system allowing detection of antibodies reactive to ancient and contemporary SARS-CoV-2 spike antigens. With stringent testing using multiple serological and neutralization assays, all the US cattle serum samples (n=598) were negative for SARS-CoV-2 antibodies. Notably, one serum sample showed borderline positive results in both pVNT and sVNT using Omicron lineage antigen. However, the sample was negative in SARS-CoV-2 live virus neutralization assay.

Although pVNTs yield comparable neutralization titer as that of live virus neutralization assays for detecting the SARS-CoV-2 antibodies[44-46], their use as a diagnostic tool could be limited due to the highly sensitive luciferase reporter system. pVNTs are widely used to determine the variant specific neutralization titer and generate antigen cartography to assess the relationship between the variants and serum antibodies[26]. We found that high percent inhibition of pseudovirus in a single serum dilution did not predict antibody detection ability using other methods, including a validated indirect ELISA. When pVNTs are repurposed to use for diagnosis using a single serum dilution, several factors may contribute to false positive results. In general, when the serum has a good titer of neutralizing antibodies to SARS-CoV-2, the percent inhibition in pVNTs are ∼100% in several two-fold serial dilutions, i.e. may be up to 1:120 dilution of serum samples, that we have observed in cat [26] and white-tailed deer[27]. Meanwhile, the cattle serum samples that were tested in pVNT showed inhibition values from 0-80% and a few samples had >80% inhibition at 1:30 dilution. Here, the reduction in pseudovirus readout could be due to cytotoxicity at 1:30 dilution, as the reduced cell growth could result in less luminescence. A way to prevent false positive results due to less cell growth is by quantitating protein concentrations in the replicate wells.

The GenScript c-Pass sVNTs are widely used for serological surveillance in humans that employ the 30% inhibition as a positive-negative cut-off[24]. SARS-CoV-2 Delta-RBD based sVNT showed 99.93% specificity and 95–100% sensitivity detecting the antibodies in humans[24]. Being a species agnostic test, sVNT has been evaluated for the antibody detection in different species including white-tailed deer, cat, hamster[47]. We have used 30% cut-off for cattle sera analysis; however, a recent study recommended the cut-off of 43% and 51% using limited numbers of pre-pandemic cattle and horse sera suggesting the cut-off of 30% may result in false positives[48]. Therefore, the Delta and Omicron-sVNT positive samples (n=3) determined in this study could be due to incorrect cut-off. This is further explained by two of the pre-pandemic samples showing more than 30% inhibition in Delta sVNT. In the in-house indirect ELISA, we have established the cut-off (mean absorabance+5 standard deviation) using 40 pre-pandemic serum samples[25]; however, this cut-off was not evaluated with the clinical samples from natural SARS-CoV-2 infection due to lack of positive samples. Therefore, we assume the possibility of false-positive results in the indirect ELISA and tested all the samples with >60% inhibition in pVNT using live virus neutralization assay. However, none of the serum samples with >60% inhibition in Delta and Omicron-pVNTs showed the neutralization of SARS-CoV-2 in live virus neutralization assays. Although, live virus neutralization assays are gold-standard comparative tests for antibody diagnostics; it requires biosafety level-3 containment facility (BSL-3)[49]. In most cases, BSL-3 laboratories are shared facilities for multiple users and requires use of expensive personnel protective equipment; therefore, it is not feasible to test large number of samples for the diagnostic purposes using SARS-CoV-2 live virus neutralization assays.

The list of susceptible animal species to SARS-CoV-2 continues to grow. Computational predictions indicate 17 bat species, and 76 rodent species have high probabilities of zoonotic capacity for SARS-CoV-2 infection[50]. Given the diversity of non-human mammalian species susceptible to SARS-CoV-2, it is possible that variants capable of infecting cattle may emerge. However, surveillance efforts in domestic and wild animal species remain inadequate to assess the spill over into animals. Our study underscores the necessity for cautious interpretation of serological diagnoses derived from these assays. Consequently, there is an urgent need to advance the development of more dependable serological assays specifically tailored to detect SARS-CoV-2 antibodies for high-risk animal populations. This emphasizes the critical importance of rigorous evaluation protocols when implementing serological assays for SARS-CoV-2 detection, thereby enhancing our ability to monitor and manage potential zoonotic transmission events.

## Acknowledgments

The study is funded by the USDA-NIFA grant (# 2020-67015-32175) (MSN and SVK), Endowed chair funds of the Penn State Huck Institutes of the Life Sciences (SVK), USDA-NIFA grant (#2023-70432-41334) (SVK and SR) and grant from Commonwealth of Pennsylvania-Department of Agriculture (SVK and SR). The authors thank Rhiannon Barry, Michele Yon, Erik Nguyen, and Manju Yadhav of the Animal Diagnostic Laboratory at Penn State, for their help in procuring the cattle serum samples.

